# Survival and Developmental Progression of Unselected Thymocytes in the Absence of the T Cell Adaptor Gads

**DOI:** 10.1101/2023.07.05.547801

**Authors:** Rose Shalah, Manal Marzouk, Enas Hallumi, Naama Klopstock, Deborah Yablonski

**Author notes:** Correspondence: Deborah Yablonski.

## Abstract

Thymic development is tightly regulated by TCR signaling via the immune adaptors SLP-76 and LAT. Gads bridges the TCR-induced recruitment of SLP-76 to LAT; yet is not absolutely required for thymic progression. To better identify Gads-dependent developmental transitions, we performed tamoxifen-induced ablation of Gads (Gads^iKO^), accompanied by expression of tdTomato, and compared the development of Gads-expressing (Tom^−^) and - ablated (Tom^+^) thymocytes within the same mouse. The frequency of Gads^iKO^ (Tom^+^) thymocytes decreased at β-selection and positive selection, confirming the Gads-dependence of these junctures; nevertheless, we observed small populations of positively-selected Gads^iKO^ SP thymocytes. Consistent with a signaling defect, expression of CD5 was strongly impaired at the β-selection checkpoint and within the DP compartment; moreover, Gads^iKO^ DP thymocytes exhibited reduced TCR-induced calcium flux. Surprisingly, MHC-non-responding (CD5^−^) Gads^iKO^ DP thymocytes exhibited reduced death by neglect; instead, aberrant populations of CD5^−^ Gads^iKO^ thymocytes progressed as far as the CD4 SP compartment, while lacking key characteristics of positively selected thymocytes. In an experimentally-induced model of death by neglect triggered by CD8 crosslinking, Gads was required for the preferential apoptosis of CD5^lo^ DP thymocytes. Our results suggest that Gads promotes passage through TCR-driven developmental checkpoints while also promoting the death by neglect of unselected thymocytes.

## Introduction

At every stage of their life cycle, T cell development, differentiation and function are tightly controlled by signaling through the clonotypic T cell antigen receptor (TCR). This principle is most clearly exemplified by thymocyte development, in which the strength of TCR signaling controls multiple developmental checkpoints (1–4).

Conventional thymocyte development proceeds through three main stages, known as double negative (DN, CD4^−^CD8^−^), double positive (DP, CD4^+^CD8^+^), and single positive (SP, either CD4^+^ or CD8^+^). The DN compartment may be further divided into four substages, based on the expression of surface markers, including DN1 (CD44^+^CD25^−^), DN2 (CD44^+^CD25^+^), DN3 (CD44^−^CD25^+^), and DN4 (CD44^−^CD25^−^) (5).

As thymocytes pass through the DN compartment, productive rearrangement of TCRβ is required for expression of the pre-TCR. Signals emanating from the pre-TCR trigger β-selection, which is characterized by rapid thymocyte proliferation and upregulation of CD5, along with passage of DN3 thymocytes through DN4 and into the DP compartment (6–8).

The mature αβ TCR is first expressed during the DP stage, where its ability to recognize self peptide-MHC (p-MHC) can be assessed by following activation markers, such as CD5 and surface TCRβ (8, 9). Low expression of these markers identifies DP cells that fail to recognize self-p-MHC (9); these cells may be removed through a process called “death by neglect” (10, 11). Evidence suggests that 80-90% of DP thymocytes may undergo death by neglect (10–12), yet the precise mechanisms triggering this process are still unknown. On the other hand, strong recognition of self-p-MHC triggers death by negative selection.

Between these two extremes, weak self-recognition triggers positive selection and transition to the SP compartments. Positively-selected SP cells are typically TCRβ^hi^ and CD5^hi^, and follow a maturation trajectory, during which the expression of CCR7 transiently increases and CD24 decreases, prior to their exit from the thymus as naive peripheral T cells. Taken together, the ability of T cells to correctly interpret the strength of TCR ligation profoundly shapes thymocyte development, eventually allowing T cells to distinguish between self and foreign antigens (1–4).

TCR ligation triggers a cascade of tyrosine kinases (13, 14), initiated by a Src-family kinase, Lck, which activates a Syk-family kinase, ZAP-70. ZAP-70 phosphorylates two adaptor proteins, LAT and SLP-76 (15–21), triggering the SH2-mediated binding of multiple signaling proteins to the adaptors (reviewed in 22, 23).

Gads is an evolutionarily conserved, Grb2-family adaptor, expressed mainly in T cells and mast cells, and required for optimal antigen receptor signaling in these cell types (24). Like Grb2, Gads consists of a central SH2 domain flanked by two SH3 domains, but also includes a unique unstructured linker. Gads C-SH3 domain binds constitutively to SLP-76 (25–27), whereas its SH2 domain mediates the cooperative binding of Gads to two LAT phospho-sites, pY171 and pY191 (18, 26, 28). Through these interactions, Gads bridges the TCR-induced recruitment of SLP-76 and its associated signaling proteins to phospho-LAT (29–33), where adaptor-associated enzymes trigger downstream responses (22, 24, 34). Among the most important and best-understood downstream pathways, SLP-76-associated Itk phosphorylates LAT-associated PLC-γ1, triggering the production of second messengers that increase intracellular calcium.

Since the best understood function of Gads is to bridge the TCR-induced recruitment of SLP-76 to LAT, it is surprising that germline deletion of these adaptors results in distinct phenotypes. Both SLP-76 and LAT are essential for thymocyte development beyond the DN3 stage (35–37). In contrast, Gads-deficient mice exhibit incomplete blocks at multiple thymic checkpoints, associated with a marked reduction in thymic size; nevertheless, the presence of prominent peripheral CD4 and CD8 T cell populations demonstrates that T cell development is at least partially Gads-independent (38, 39). To date, the precise thymic developmental transitions controlled by Gads are not fully understood. We hypothesized that Gads fine tunes TCR responsiveness and thereby may affect multiple developmental checkpoints. To test our hypothesis, we established a genetic model that enabled us to compare the development of Gads-expressing and Gads-ablated thymocytes, side by side in the same mouse.

This approach revealed multiple points at which Gads influences thymocyte development. Gads was required for efficient β-selection and for optimal p-MHC-induced expression of CD5, an important correlate of positive selection. On the other hand, Gads was required for efficient death by neglect of MHC-non-responsive DP thymocytes; in its absence, unselected, CD5^−^ thymocytes progressed as far as the CD4 SP compartment. This observation sheds an unexpected light on the involvement of TCR signaling elements in the regulation of death by neglect.

## Materials and Methods

### Antibodies and staining reagents

Anti-mouse antibodies CD4-PerCP-Cy5.5 or -FITC (Clone RM4-4 and GK1.5), CD8-APC-Cy7, or –APC, or –FITC (Clone 53-6.7 or QA17A07), CD44-BV421 (Clone IM7), CD5-BV711 (Clone 53-7.3), CD25-APC (Clone PC61), NK1.1-FITC (Clone PK136), TCR γδ-FITC (Clone GL3), TCR β-BV605 (Clone H57-597), CCR7-AF488 (Clone 4B12), Ly-6G/Ly-6C (Gr-1)-FITC (clone RB6-8C5), CD45R/B220-FITC (clone RA3-6B2), TER-119-FITC (clone TER-119), CD11b (Mac1)-Ly40-FITC (clone M1/70), Zombie Aqua Fixable viability kit, and Biotin-conjugated anti-mouse CD3ε (clone145-2C11), CD4 (clone RM4-4) and CD8 (clone 53-6.7) were from Biolegend. Cleaved caspase 3-AF647 (Clone D3E9) was from Cell Signaling. Gads-AF647 (UW40) was from Santa Cruz Biotechnology. Unless noted otherwise, all PBS used in this study is without calcium and magnesium.

### Mice

Mice were on the C57BL/6 background and were maintained in an SPF facility under veterinary supervision and in accordance with institutional ethics guidelines. Gads-deficient (Gads^−/−^) mice (38) were generously provided by C. Jane McGlade (University of Toronto). Gads^FL^ mice, in which LoxP signals flank Grap2 (Gads) exon2, were generated for us by Cyagen/Taconic using a CRISPR/Cas9 approach. Gads^FL^ mice were cross-bred to Jackson Laboratory strain #007914 (Ai14), to incorporate a Cre-dependent tdTomato (Tom^FL-stop^) reporter at the Rosa26 locus (40). The ubiquitously-expressed, tamoxifen-inducible UBC-Cre-ERT2 Cre recombinase (41) was maintained in a hemizygous state. To generate mice for experiments, Gads^FL/FL^Tom^FL-stop/FL-stop^ mice were bred to Gads-deficient Cre driver mice (Gads^−/−^UBC-Cre-ERT2^+^), to generate Gads^FL/−^Tom^FL-stop^ progeny, either Cre^+^ or Cre^−^.

### Tamoxifen-inducible deletion of Gads

Tamoxifen (Sigma, T5648) was dissolved at 10 mg/ml in fresh corn oil by shaking at 37°C for 3-4 hr. For tamoxifen-inducible deletion of Gads, 6-7 week old Gads^FL/−^Tom^FL-stop^ mice, either UBC-Cre-ERT2^+^ or Cre^−^, were treated with tamoxifen daily for five days by intraperitoneal injection (75 μg/g mouse weight). To assess the efficiency of Cre-mediated recombination, 200 μl of blood was collected from the cheek into 1.5ml Eppendorf tubes containing 50 units heparin. Blood samples were diluted to 2ml in PBS and underlayed with 2 ml of Histopaque-1083 (Sigma, 10831). Following centrifugation at 400 g for 30 min at room temperature, leukocytes were collected from the interface, and washed once in PBS prior to staining for FACS analysis.

### Thymocyte isolation and staining

A single cell suspension of thymocytes was generated by pressing the thymus through a 40 μm Cell Strainer. Cells were washed once with PBS supplemented with 2% FCS, then washed and re-suspended in 10 ml PBS without FCS. Cells were stained for 30 min at RT in the dark with Zombie Aqua Fixable Viability Kit. Staining was terminated by washing cells with 2ml FACS buffer (PBS, supplemented with 2% FCS and 0.02% sodium azide). Cells were then stained for 15-30 min at 4°C with a cocktail of fluorescently labeled antibodies in FACS buffer, washed once with FACS buffer and once with PBS, fixed for 20 min at RT in the dark with a 1:1 dilution of fixation buffer in PBS (IC Fixation Buffer Thermo Fischer Scientific, 00-8222-49), then washed in FACS buffer. For intracellular staining with Gads, TCR or cleaved caspase 3, fixed cells were washed twice with 1x Thermo Fischer Scientific permeabilization buffer (Invitrogen, 00-8333-56), prior to intracellular staining in the same buffer.

### Annexin V staining

After staining the cells with zombie and with the desired surface markers, cells were washed with Annexin V Binding Buffer (Biolegend, 422201) and stained for 15 min at RT with annexin V (Biolegend, 640919).

### CD8-induced death by neglect

Freshly-isolated thymocytes were incubated at 37°C for 15 min in Hepes-buffered complete T cell medium (high glucose DMEM, supplemented with 10% iron fortified bovine calf serum, NEAA (non-essential amino acids), 2mM glutamax, 1mM sodium pyruvate, 50 μM β-mercaptoethanol, pen-strep, and 20mM Hepes, pH7.3). Avidin beads (Spherotech, SVP-30-5) were pre-incubated in the same medium with different concentrations of CD4- or CD8-biotin and washed once. To induce death, 0.25X10^6^ cells were mixed in a round-bottomed well with 1.25X10^6^ pre-coated avidin beads. After incubation at 37°C for 1 hour, the cells were washed once with cold FACS buffer, stained at 4°C with surface antibodies for CD4, CD8 and CD5, washed once with FACS buffer and once with annexin V binding buffer, and stained for 15 min at RT with annexin V and DAPI. To calculate the induced death index, the net CD8-induced increase in the early apoptotic population was expressed as a fraction of the population that was live at baseline ((%AV^+^DAPI^−^_exp_-%AV^+^DAPI^−^_baseline_)/ %AV^−^DAPI^−^_baseline_).

### FACS analysis

Stained cells were read on a Fortessa II FACS analyzer, using single-stained cells or compensation beads as controls. Results were analyzed with FlowJo™ software, while gating on live (Zombie-negative) single cells. Where indicated, a FITC-labeled dump gate was used to exclude NK1.1^+^ and γδ^+^ T cells, as well as cells that stained positive for Ly-6G/Ly-6C (Gr-1), CD45R/B220, TER-119, or CD11b (Mac1)-Ly40.

### Calcium fluorimetry

Calcium flux was measured using a mixture of thymocytes from tamoxifen-treated Cre^+^ and Cre^−^ mice. Thymocytes were washed with calcium staining buffer (RPMI supplemented with 1% FCS, 20mM Hepes pH 7.3, and 2mM glutamine), and incubated in the dark for 30 min at 30°C, with 0.5 ml per each 5 million cells of calcium staining buffer containing 4μM indo-1 AM (eBioscience), 2mM probenecid, fluorescent cell surface markers, including CD4-FITC (clone GK1.5) and CD8-APC (clone QA17A07), and biotinylated stimulatory antibodies: CD3, CD4 (clone RM4-4) and CD8 (clone 53-6.7), 3μg/ml each. The stained cells were then washed twice and resuspended in RPMI supplemented 20mM Hepes pH 7.3, and 2mM Glutamine. Cells were stored on ice and preheated to 37°C for 5 min prior to ratiometric calcium fluorimetry, which was performed by FACS, with the temperature maintained at 37 °C, and TCR stimulation was induced by adding 25 μg/ml streptavidin (Biolegend, 405150) at the 60 s time point. Excitation was at 355 nm and emission was measured using the bandpass filters 405/20 (Ca-bound indo-1) and 485/22 (free indo-1), with data presented as the ratio between these two values. Data analysis was performed while gating on CD5^+^, Tom^+^ or Tom^−^ DP thymocytes.

### Statistical Analysis

All experiments in this study were performed two or more times, with similar results. Statistical analysis was performed using the Prism software package. p values were determined using the statistical tests indicated in figure legends. The following symbols were used to indicate statistically significant p values: *, p<0.05; **, p<0.01; ***, p<0.0005; ****, p<0.0001.

## Results

### A mouse model for inducible deletion of Gads, accompanied by the expression of tdTomato

To better distinguish Gads-dependent and -independent developmental pathways, we developed a mouse model system for the inducible deletion of Gads. This model is comprised of two “floxed” genes, Gads^FL^ and Tom^FL-stop^ (Fig 1A), controlled by a ubiquitously-expressed tamoxifen-inducible Cre recombinase, UBC-Cre-ERT2 (41). Cre-mediated removal of Gads exon 2 creates a frameshift mutation that alters the protein sequence after K26, resulting in a premature stop codon after 29 amino acids (Fig 1B). Tom^FL-stop^ is a previously-described fluorescent reporter, expression of which is blocked by a LoxP-flanked, upstream, in-frame stop codon (40). Cre-mediated removal of this stop cassette is required for expression of the tdTomato gene product. Together, this system permits Cre-mediated deletion of Gads, accompanied by the expression of tdTomato.

**Fig 1.**
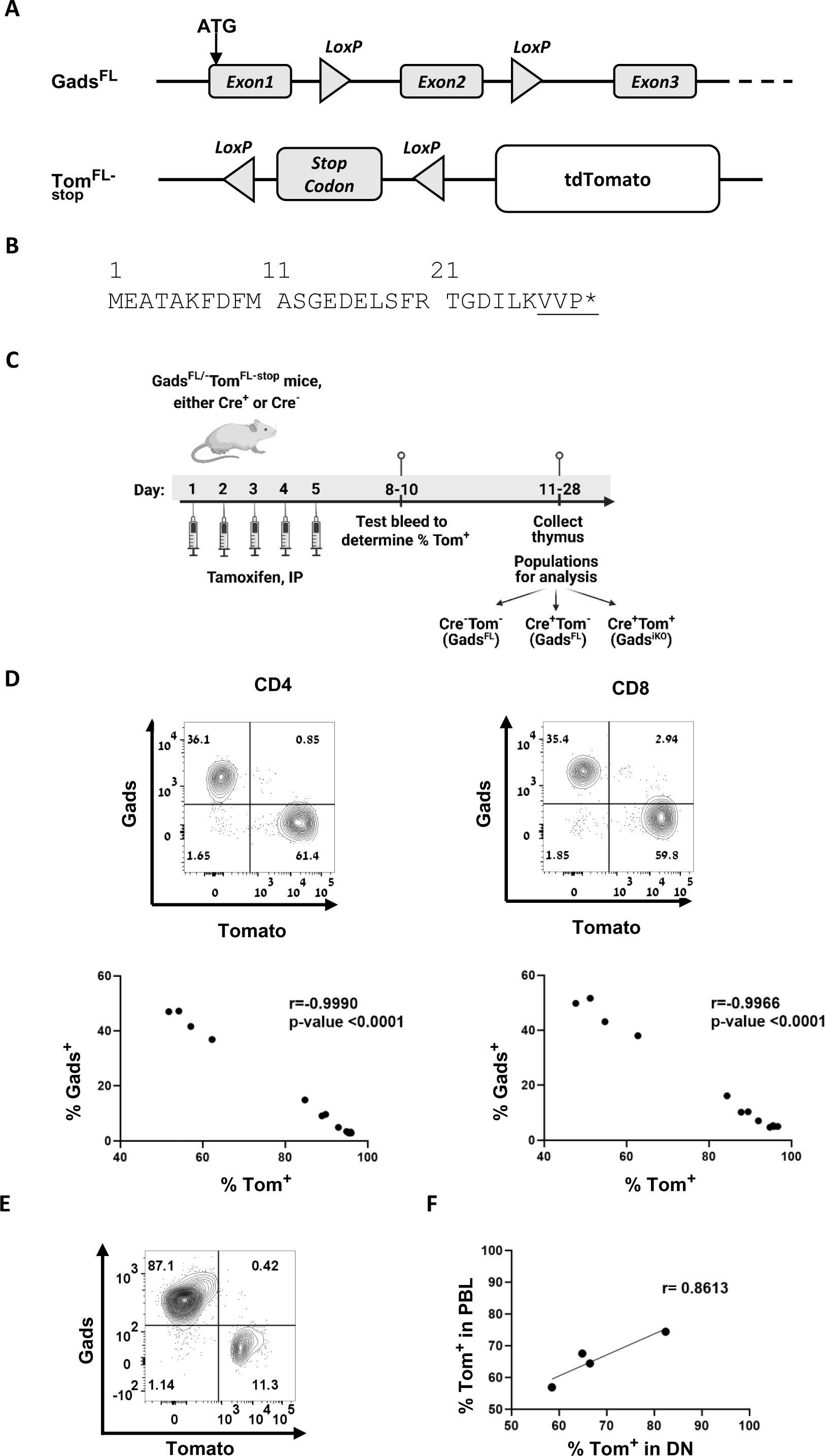
A mouse model for inducible deletion of Gads, accompanied by the expression of tdTomato. The Gads^iKO^ mouse model comprises two “floxed” genes, whose expression is controlled by UBC-Cre-ERT2, a ubiquitously expressed tamoxifen-inducible Cre recombinase (41). (**A**) Prior to Cre activity, LoxP sites flank exon 2 of *Grap2*, the gene that encodes Gads. Within the *Rosa26* locus, a second set of LoxP sites flank an in frame stop codon that prevents the expression of a fluorescent marker, tdTomato (40). (**B**) Cre-mediated recombination removes exon 2 of *Grap2*, creating a frameshift mutation that alters the protein sequence after K26 and results in a premature stop codon after 29 amino acids. (**C**) Experimental workflow for tamoxifen-inducible ablation of Gads. Created with BioRender.com. (**D**) One week after starting tamoxifen treatment, peripheral blood leukocytes (PBL) were stained for surface markers and intracellularly for Gads. Top: representative results obtained while gating on CD4^+^ or CD8^+^ T cells. Bottom: The frequency of Gads^+^ and Tom^+^ phenotypes observed in the CD4^+^ and CD8^+^ peripheral blood populations of 13 individual Cre^+^ mice. The Pearson correlation coefficient (r) was calculated using Prism. (**E**) Two weeks after the start of tamoxifen, thymocytes were stained intracellularly for Gads. Representative result is presented while gating on live thymocytes. (**F**) Two weeks after the start of tamoxifen, total thymocytes were stained to identify DN (CD4^−^CD8^−^) cells. The frequency of Tom^+^ DN thymocytes correlated with the frequency of Tom^+^ PBLs observed four days earlier in the same mice.

### The tdTomato marker reliably identifies Gads^iKO^ peripheral T cells and thymocytes

Our typical experimental workflow is depicted in Fig 1C. For inducible deletion of Gads, we administer tamoxifen for five consecutive days to UBC-Cre-ERT2^+^ Gads^FL/−^Tom^FL-stop^ mice; in parallel, we tamoxifen-treat an equal number of control Cre^−^ Gads^FL/−^Tom^FL-stop^ litter-mates. In this configuration, Cre-mediated recombination at a single allele of Gads and a single allele of tdTomato suffices to render the cells tdTomato-positive and Gads-deficient (Tom^+^Gads^iKO^).

Tom^+^ peripheral blood T cells were reproducibly observed one week after the initiation of tamoxifen treatment; however, their frequency varied between mice. The rapid appearance of Tom^+^ peripheral T cells suggests that they derive from Cre activity in pre-existing peripheral populations.

To more directly detect ablation of Gads, we stained PBLs intracellularly with a fluorescently-labeled anti-Gads antibody, and analyzed the results while gating on CD4^+^ or CD8^+^ T cells. Tom^+^ peripheral T cells were uniformly Gads-negative, (Gads^iKO^, Fig 1D, top panels). A tight correlation between the Tom^+^ and Gads^iKO^ phenotypes was apparent as soon as one week after the initiation of tamoxifen treatment (Fig 1D, bottom panels) and remained intact as long as 5 weeks after the initiation of tamoxifen treatment (data not shown). Within the thymus, Tom^+^ cells were likewise uniformly Gads-negative (Fig 1E). These results establish Tom^+^ as a marker that reliably identifies Gads^iKO^ cells in our experimental system.

Thymocyte development begins in the DN quadrant, which is first populated by bone-marrow-derived common lymphoid progenitors (42). Two weeks after the initiation of tamoxifen treatment, the frequency of Tom^+^ DN thymocytes strongly resembled the frequency of Tom^+^ PBLs in the same mouse (Fig 1F). This observation suggests that tamoxifen triggered UBC-Cre-ERT2-mediated recombination with comparable efficiency in thymic progenitor cells and in preexisting peripheral T cell compartments.

### Prior to Cre-mediated ablation, Gads^FL^ supports wild-type functionality

To validate the inducible effect of our model, we first verified that, prior to its Cre-mediated ablation, Gads^FL^ supports wild type functionality. To this end, we compared thymic development in mice of three genotypes: Gads^+^ (wild-type), Gads^FL^, and Gads^−^ (germ-line Gads-deficient mice).

As previously reported (38, 39), Gads^−/−^ mice had markedly reduced thymic cellularity (Fig 2A), and reduced frequency of CD4 SP thymocytes (Fig 2B), accompanied by a reduced ratio of CD4 to CD8 SP thymocytes (Fig 2C). A partial block at the DN to DP transition was reflected in an increased frequency of DN thymocytes, and reduced frequency of DP thymocytes (Fig 2D-F). In all of these parameters, we observed statistically significant differences between Gads-expressing and Gads-deficient thymocytes, but no significant differences were observed between thymocytes expressing wild type and floxed Gads (Fig 2A-C and E-F).

**Fig 2.**
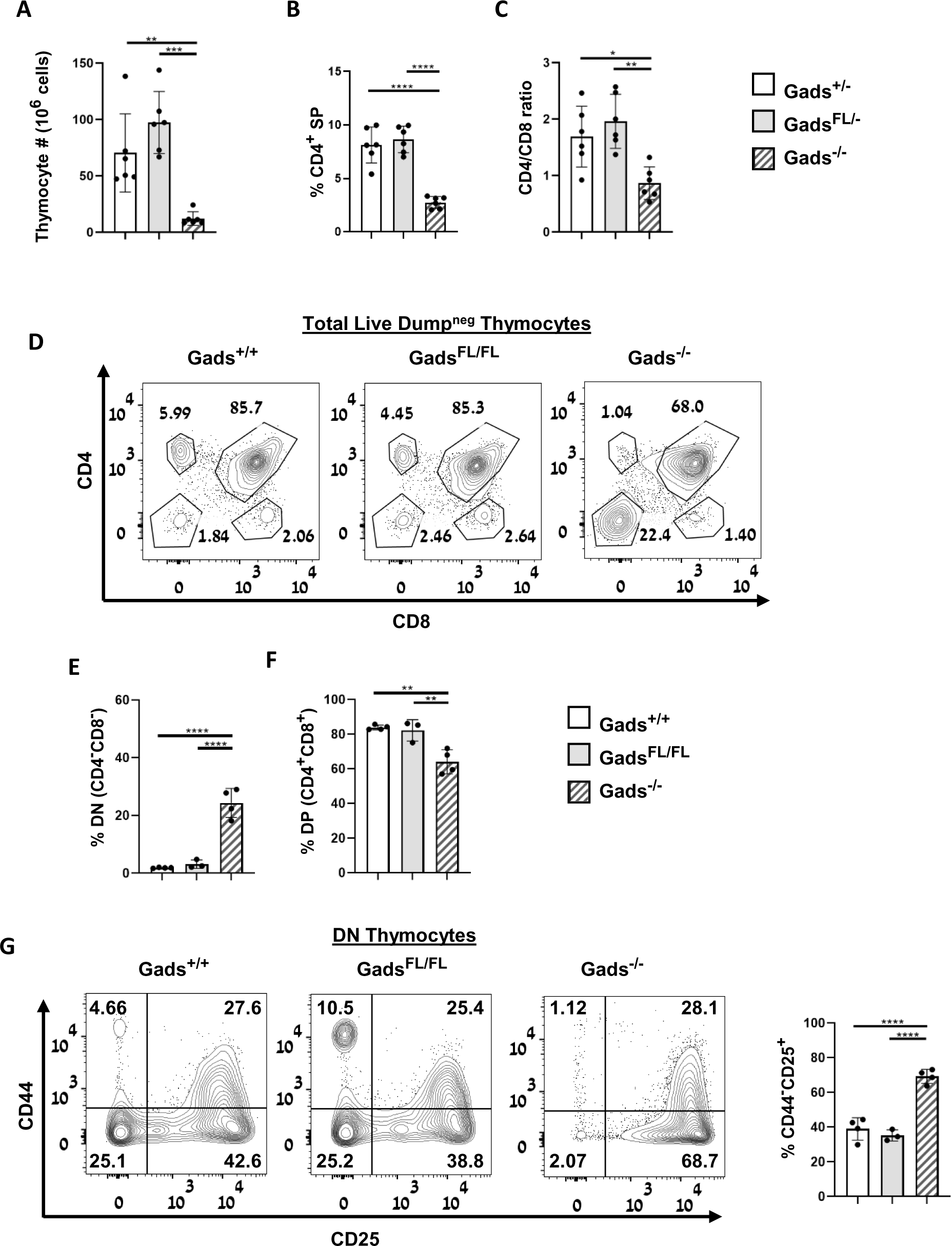
Prior to Cre-mediated deletion, Gads^FL^ thymocytes exhibit wild type phenotypes. FACS analysis of thymocytes from 7-8 week old mice. None of the mice in this analysis expressed Cre. (**A-C**) Mice were Gads^+/−^, open bars; Gads^FL/−^, grey bars; Gads^−/−^, striped bars. n=6 mice for each genotype. (**A**) Thymic cellularity. (**B**) The frequency of CD4 SP thymocytes. (**C**) The ratio of CD4 SP to CD8 SP thymocytes. (**D-G**) To facilitate quantitative analysis of the DN compartment, a mixture of FITC labeled antibodies was used to create a “dump gate” to exclude NK1.1^+^ and TCRγδ^+^ thymocytes, as well as erythrocyte, granulocyte, macrophage, monocyte and B cell lineages. Mice were Gads^+/+^, open bars; Gads^FL/FL^, grey bars; Gads^−/−^, striped bars. n=3-4 mice for each genotype. (**D**) Representative result depicts CD4 and CD8 expression, while gating on live (zombie-negative), dump^neg^ thymocytes. (**E-F**) The percent of thymocytes found within each of the indicated gates, as defined in D. (**G**) To assess progression through DN substages, the expression of CD44 and CD25 was measured while gating on live (zombie-negative), dump^neg^ DN thymocytes. Left: representative result. Right: the percent of DN thymocytes found in DN3 (CD44^−^CD25^+^). Error bars indicate the standard deviation. Statistical significance was determined by one-way ANOVA.

Within the DN compartment, Gads^−/−^ thymocytes exhibited the previously reported partial block at the DN3 to DN4 transition (38, 39). In this trait as well, Gads^−/−^ DN thymocytes exhibited a statistically significant increase in the frequency of DN3 cells, which was not observed in Gads^FL/FL^ mice (Fig 2G). These observations provide strong evidence that the LoxP sites surrounding Gads exon 2 do not impair the expression or function of Gads.

### Cell-autonomous impairment of β-selection in Gads^iKO^ (Tom^+^) thymocytes

Following tamoxifen treatment, the contemporaneous development of Tom^+^ (Gads^iKO^) and Tom^−^ (Gads-expressing) thymocytes within the same mouse provided an opportunity to assess the cell-autonomous effects of Gads on thymocyte development.

To focus on conventional T cell development, we used a “dump” gate to exclude NK1.1^+^ and TCRγδ^+^ thymocytes as well as non-T lineage cells. We followed thymic progression within the dump^neg^ population, while gating separately on Gads-expressing (Tom^−^) and Gads^iKO^ (Tom^+^) thymocytes.

Tom^+^ (Gads^iKO^) thymocytes largely phenocopied the germline Gads-deficient mouse, exhibiting a partial block at the DN to DP transition compared to Cre^−^ thymocytes (Fig 3A). This developmental block was reflected in a marked, statistically-significant increase in the frequency of DN thymocytes (Fig 3B) and reduced frequency of CD4 SP thymocytes (Fig 3C). The frequency of CD8 thymocytes was unchanged (Fig 3D) resulting in a reduced CD4/CD8 ratio (Fig E). In contrast, Tom^−^ thymocytes from the same Cre^+^ mice closely resembled Cre^−^ thymocytes (Fig 3A-E).

**Fig 3.**
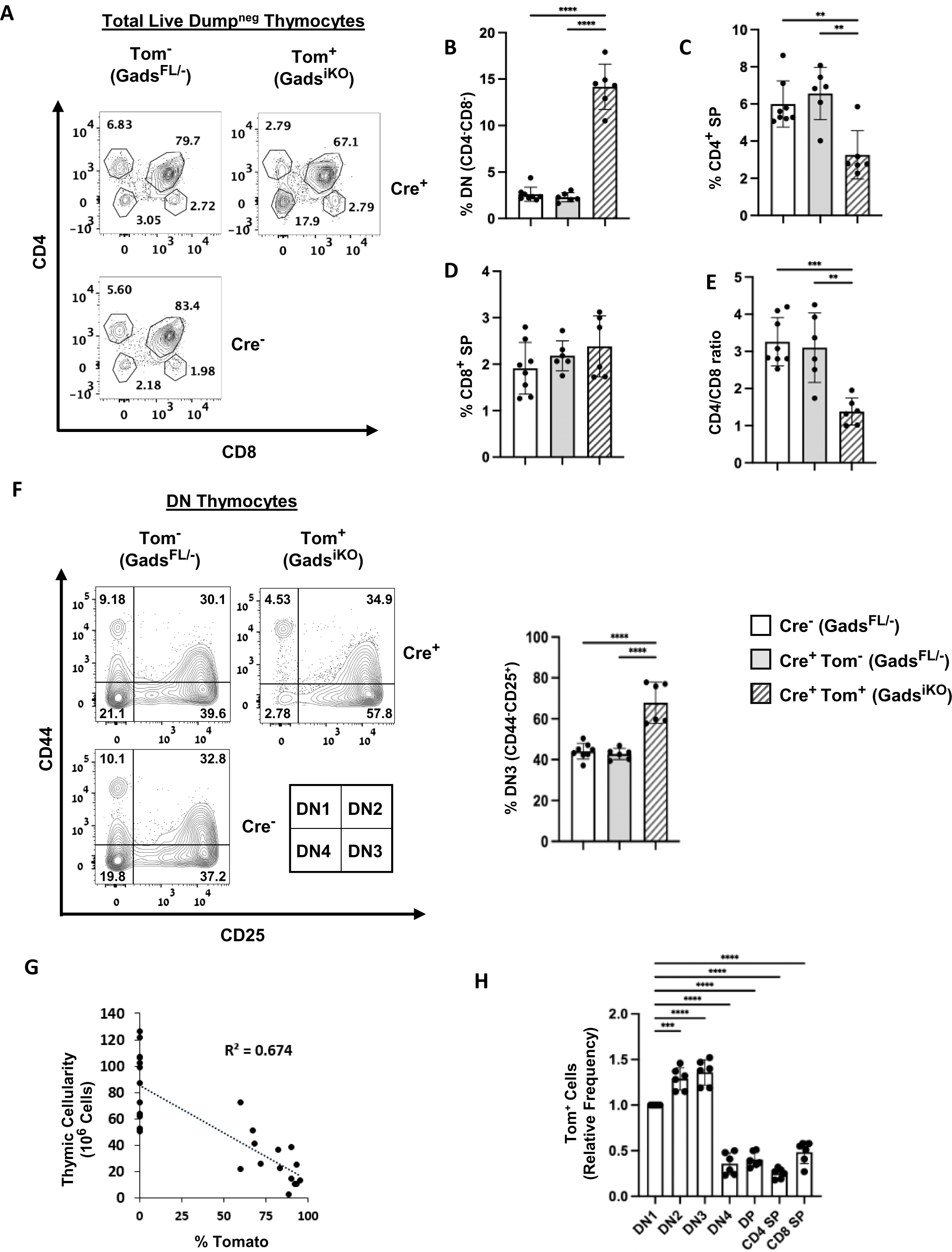
Cell autonomous developmental phenotypes of Gads^iKO^ thymocytes. Mice were tamoxifen treated as in Fig 2A, and thymi were collected 2.5 weeks after starting tamoxifen. (**A-F and H**) Thymic developmental markers were assessed by FACS using a cohort of mice in which 60-75% of Cre^+^ PBLs were Tom^+^. Zombie staining was used to exclude dead thymocytes. A mixture of FITC labeled antibodies was used to create a “dump gate” to exclude NK1.1^+^ and TCRγδ^+^ thymocytes, as well as erythrocyte, granulocyte, macrophage, monocyte and B cell lineages. (**A**) Representative distribution of CD4 and CD8 surface markers within the live, dump^neg^ population, assessed while gating separately on Tom^−^ (Gads^FL/−^) and Tom^+^ (Gads^iKO^) thymocytes. (**B-D**) The percent of thymocytes found within each of the subpopulations defined in A. (**E**) The ratio of CD4 SP to CD8 SP thymocytes. (**F**) Progression through the DN substages was assessed while gating separately on live, dump^neg^ Tom^+^ and Tom^−^ DN (CD4^−^CD8^−^) thymocytes. Left: Representative result. Right: Quantitation of the results using the gating strategy shown at left. The legend relates to bar graphs B-F: Cre^−^ (Gads^FL/−^), open bars; Cre^+^Tom^−^ (Gads^FL/−^), grey bars; Cre^+^Tom^+^ (Gads^iKO^), striped bars. n = 8 Cre^−^ and 6 Cre^+^ mice. Error bars indicate the standard deviation. Statistical significance was determined by a one-way ANOVA test, Tukey’s multiple comparison test. (**G**) Reduced thymic cellularity in Gads^iKO^ mice. Total thymic cellularity is plotted as a function of the frequency of Tom^+^ PBLs. Cre^−^ mice are plotted on the Y axis. A linear regression confirmed a tendency to reduced thymus size with increased frequency of Tom^+^ cells. This panel combines data from multiple experiments in which the thymus was collected 1.5-2.5 weeks after starting tamoxifen. n= 13 Cre^−^ and 14 Cre^+^ mice. (**H**) The percent of Tom^+^ thymocytes was determined while gating separately on each of the indicated sub-populations. To facilitate statistically valid comparisons between mice, the frequency of Tom^+^ cells in each sub-stage was normalized to their frequency in the DN1 compartment of the same mouse. Statistical significance was determined by a one-way ANOVA test, Dunnett’s multiple comparisons test. (n=6 Cre^+^ mice).

Within the DN compartment, Gads^iKO^ thymocytes exhibited a partial block at the DN3 to DN4 transition; in this trait as well Cre^+^Tom^−^ DN thymocytes phenocopied Cre^−^ DN thymocytes, and both were statistically different from Gads^iKO^ DN thymocytes (Fig 3F).

We considered the possibility that selective impairment of Gads^iKO^ thymocytes may allow for compensatory progression of Tom^−^ thymocytes, which would crowd out their Tom^+^ counterparts. Since Gads-deficient mice are characterized by a small thymus (Fig 2A and 38, 39), we reasoned that cell-autonomous effects of Gads would result in a small thymus whereas compensatory progression of Tom^−^ thymocytes may maintain normal thymic cellularity. Cellularity was rapidly and substantially reduced in tamoxifen-treated Cre^+^ mice, relative to Cre^−^ littermates (Fig 3G). This trend was most apparent in mice with the highest frequency of Tom^+^ PBLs.

Taken together, our results suggest that the phenotypic effects of deleting Gads are predominantly cell-autonomous, with Tom^−^ thymocytes largely following normal developmental pathways, while Tom^+^ (Gads^iKO^) thymocytes within the same mouse phenocopy germ-line Gads-deficient thymocytes.

These observations suggested a simple strategy to identify developmental junctions at which Gads exerts its effects. We posited that the frequency of Tom^+^ thymocytes should decrease as cells pass through Gads-dependent developmental transitions. Conversely, the frequency of Tom^+^ thymocytes may remain the same or even increase in Gads-independent lineages. Consistent with the well-established partial blockage of Gads-deficient thymocytes at the β-selection and positive-selection checkpoints (38, 39), the frequency of Tom^+^ cells dropped substantially as cells passed from DN3 into DN4 (Fig 3H).

### Developmentally-regulated expression of CD5 depends on Gads

Thymic progression is driven by pre-TCR and TCR signaling, which are accompanied by surface expression of CD5 (8). To assess how Gads may affect TCR signaling *in situ*, we examined CD5 expression as a function of the developmental stage.

CD5 expression within the DN compartment was assessed while using a dump gate to exclude non-conventional NK1.1^+^ and TCRγδ^+^ DN thymocytes. TCRβ^−^CD5^−^ DN2 cells represent a relatively <u>early DN</u> population (Fig. 4A, top row, pop. 1), that was used for normalization in this analysis. Within DN3, TCRβ expression was associated with moderate upregulation of CD5 in Gads-expressing thymocytes, but not in Gads^iKO^ thymocytes (Fig 4A, 2^nd^ row). We refer to TCRβ^+^ DN3 cells that failed to upregulate CD5 or transit into DN4, as a population exhibiting <u>arrested β-selection</u> (Fig 4A, pop 2).

**Fig 4.**
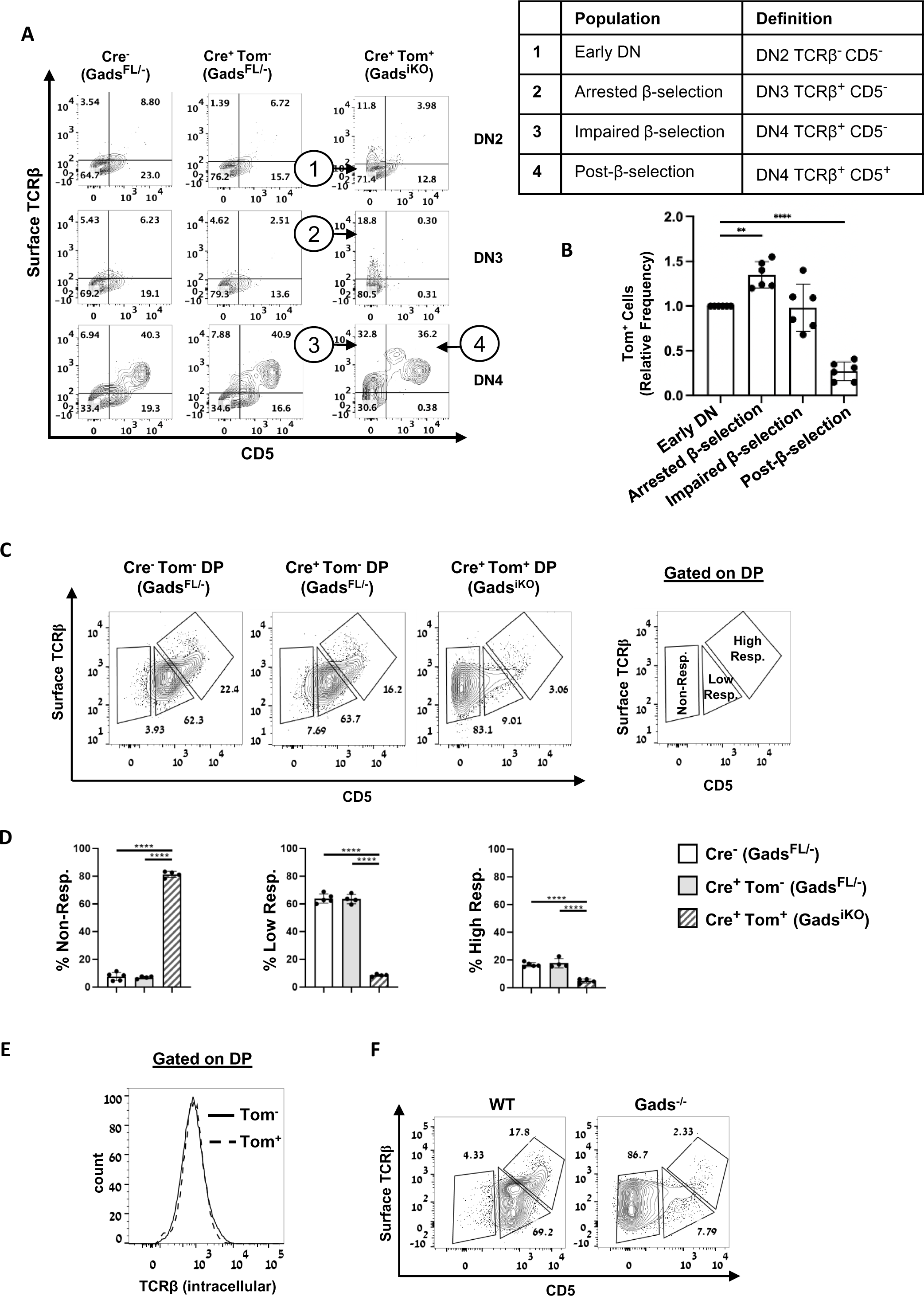
Developmentally-regulated expression of CD5 depends on Gads. Thymocytes were analyzed 2-2.5 weeks after the initiation of tamoxifen treatment. (**A-B**) CD5 expression within the DN population. Data were collected while using a dump gate to exclude DP, SP and non-conventional NK1.1^+^ and TCRγδ^+^ thymocytes as well as non-T cell lineage cells. **(A)** Representative distribution of TCRβ and CD5 expression within each DN subcompartment, as defined in Fig 3F Subpopulations of interest are indicated by numbers and defined in the table at right. (**B**) The percent of Tom^+^ cells within each of the DN subpopulations defined in A was normalized to the percent of Tom^+^ cells in the early DN (DN2 TCRβ^−^ CD5^−^) compartment of the same mouse (n = 6 Cre^+^ mice). Error bars indicate the standard deviation. Statistical significance was determined by one-way ANOVA, Dunnett’s multiple comparisons test. (**C-E**) TCR and CD5 expression within the DP population. (**C**) Representative result obtained while gating on live DP (CD4^+^CD8^+^) thymocytes. DP subpopulations are defined at right. (**D**) The percent of DP cells within each of the subpopulations defined in C. Statistical significance was determined by a one-way ANOVA Tukey’s multiple comparison test. (**E**) TCRβ expression in the DP compartment was detected by intracellular staining. (**F**) Thymocytes from 8.5-9 week old WT and conventional Gads KO mice were stained and analyzed by FACS, while gating on the DP population. Representative result.

Among DN4 thymocytes, we observed an unusual population of Gads^iKO^ cells exhibiting <u>impaired β-selection</u> (Fig 4A, pop 3), as they remained CD5^−^ despite expressing TCRβ. In contrast, Gads-expressing, TCRβ^+^ DN4 thymocytes were uniformly CD5^+^, suggesting that they represent a <u>post β-selection</u> population (Fig 4A, pop 4), composed of thymocytes that have responded to pre-TCR signaling by passing the β-selection checkpoint.

To better define the juncture at which Gads affects conventional β-selection, we examined the frequency of Tom^+^ cells in each of the above populations of interest, relative to their frequency in the early DN population. The relative frequency of Tom^+^ cells increased in the arrested β-selection population, and markedly decreased in the CD5^+^, post β-selection population (Fig 4B). Taken together, these results demonstrate that Gads is required for pre-TCR-induced expression of CD5, along with the associated progression from DN3 to DN4.

In conventional T cell development, β-selection is followed by progression to the DP compartment, where TCR-mediated recognition of self-peptide-MHC antigens induces further upregulation of CD5 and TCRβ. Investigators have estimated TCR-driven progression through the DP compartment by subdividing it according to the surface expression of CD5 and TCRβ (9, 43). In a similar fashion, we subdivided the DP compartment into non-responding (CD5^−^), low responders (CD5^med^TCRβ^med^) and high responders (CD5^hi^TCRβ^hi^) (Fig 4C). Gads^iKO^ (Tom^+^) DP thymocytes were significantly more likely to be non-responding and less likely to be low or high responders (Fig 4C and D). Indeed, a prominent, CD5^−^ Gads^iKO^ DP population was observed at all time points that we examined from 2-6.5 weeks after the initiation of tamoxifen treatment (data not shown). TCR expression, as detected by intracellular staining with TCRβ, was comparable in Gads^iKO^ and Gads-expressing DP thymocytes (Fig 4E). We therefore attribute the predominantly non-responding phenotype of Gads^iKO^ DP thymocytes to a TCR signaling defect.

The DP subset distribution was similarly skewed in germ-line Gads-deficient mice (Fig 4F), suggesting that a predominantly CD5^−^ (non-responding) DP phenotype is a previously unreported, core characteristic of Gads-deficient thymocytes.

### Profound defect in TCR signaling in Gads-deficient DP thymocytes

Despite the absence of Gads, a small population of Gads^iKO^ DP thymocytes expressed CD5 (Fig 5A), raising the possibility that a subset of Gads^iKO^ DP thymocytes may be TCR signaling-competent. To test this notion, we measured TCR-induced calcium flux, using an *ex vivo*, FACS-based assay. CD5^−^ DP cells did not increase calcium in this assay (data not shown), we therefore present data that was obtained while gating on CD5^+^ DP thymocytes. This gate produced well-matched surface TCRβ expression in Tom^+^ and Tom^−^ cells (Fig 5B, left). Moreover, intracellular staining verified that Tom^+^ DP cells within the CD5^+^ gate were indeed Gads-deficient (Fig 5B, right).

**Fig 5.**
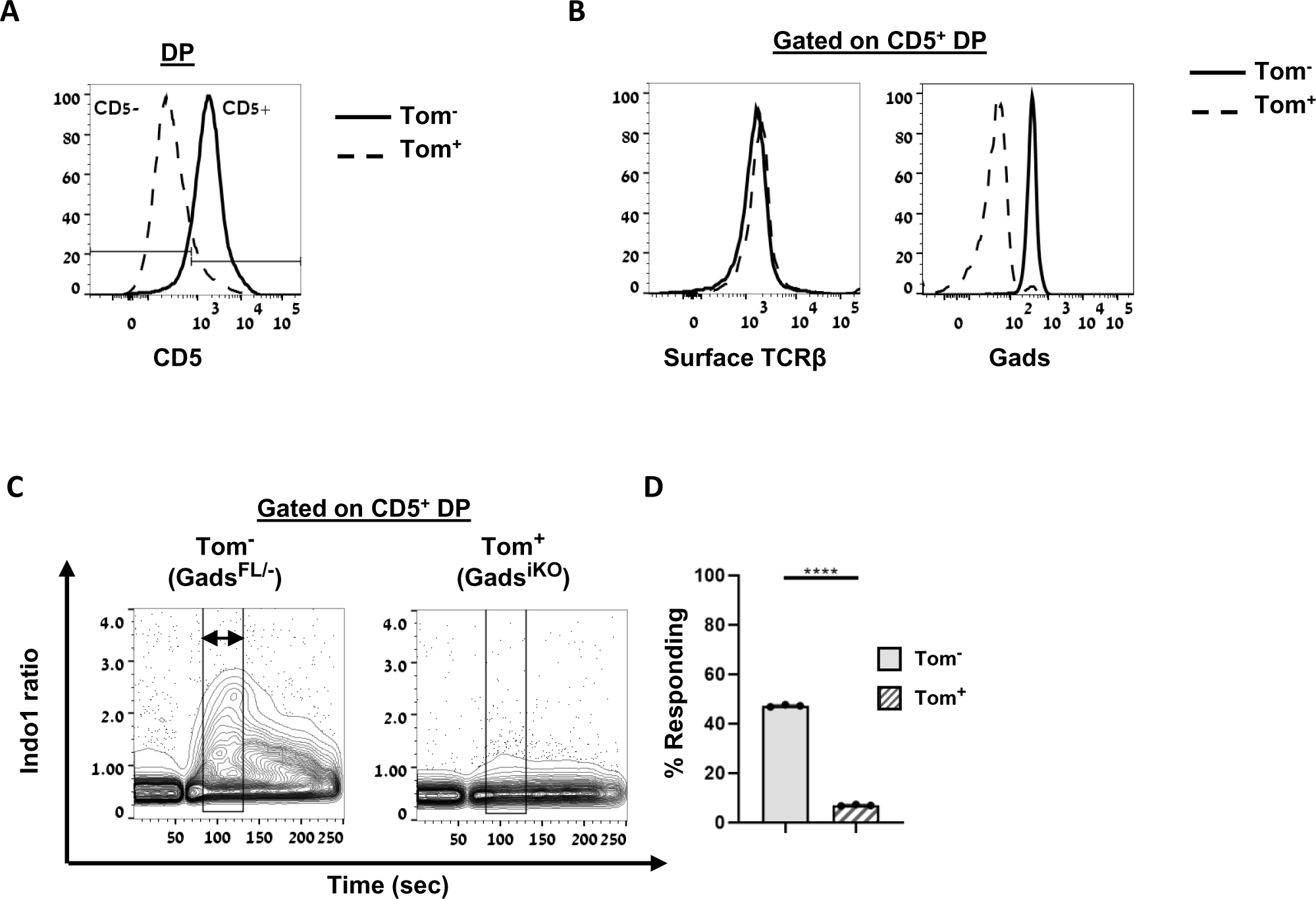
Gads promotes TCR responsiveness in the DP compartment. 2 weeks after the initiation of tamoxifen treatment, Tom^+^ and Tom^−^ thymocytes were mixed together and stained with the calcium indicator, indo1, fluorescent cell surface markers, and biotinylated stimulatory antibodies (CD3, CD4 and CD8). (**A**) CD5 expression within the DP gate. (**B**) Expression of surface TCR and intracellular Gads within the CD5^+^ gate, as defined in A. (**C-D**) Calcium fluorimetry was performed by FACS at 37°C, with stimulation initiated at the 60 sec time point by the addition of streptavidin. The indo1 ratio is shown as a function of time, while gating on CD5^+^, Tom^+^ or Tom^−^, DP thymocytes. (**C**) Representative result. The double headed arrow marks an ∼50 sec time window within which we calculated the percent of responding cells. (**D**) The percent of cells exhibiting an indo1 ratio >1 within the time window shown in D. n=3 repeats. Error bars were too small to depict. Statistical significance was determined by unpaired T-test.

To measure calcium, a mixture of Tom^+^ and Tom^−^ thymocytes was fluorescently stained with anti-CD4, -CD8 and -CD5, along with biotinylated stimulatory antibodies, and stimulation was induced by the addition of streptavidin. As a control for cell viability and intact calcium stores, ionomycin-induced calcium flux was not affected by Gads (data not shown).

After measuring baseline calcium at 37°C, cells were triggered by streptavidin-mediated co-cross-linking of CD3, CD4 and CD8, to mimic co-receptor-dependent recognition of MHC. Intracellular calcium increased in Gads-expressing (Tom^−^) DP thymocytes, but this response was blunted in Gads^iKO^ DP thymocytes (Fig 5C). When analyzed on a single-cell basis, the frequency of responding CD5^+^ DP thymocytes was reduced 6.8-fold in the absence of Gads (Fig 5D). These data confirm a profound signaling defect in Gads-deficient DP thymocytes, even within the CD5^+^, potentially responsive gate.

### Gads is required to facilitate death by neglect within the TCR-non-responding DP population

Peptide-MHC-non-responsive DP thymocytes are candidates for death by neglect (11), a multi-step, apoptotic process, in which cleavage-mediated activation of caspase 3 is followed by surface exposure of phosphatidyl serine, which can be measured by staining with annexin V. The marked accumulation of CD5^−^, non-responding Gads^iKO^ DP thymocytes raised the question of whether Gads may influence the process of death by neglect.

The average frequency of cleaved caspase 3^+^ cells within the non-responding DP gate was slightly reduced in the absence of Gads (Fig 6A), but this difference was not statistically significant, due to wide inter-mouse variation (data not shown). A statistically significant trend was observed upon pairwise comparison of Tom^+^ and Tom^−^ non-responding DP thymocytes within the same mouse (Fig 6B). Further downstream, the frequency of annexin V^+^ non-responding DP cells was substantially and significantly reduced in the absence of Gads (Fig 6C-D).

**Fig 6.**
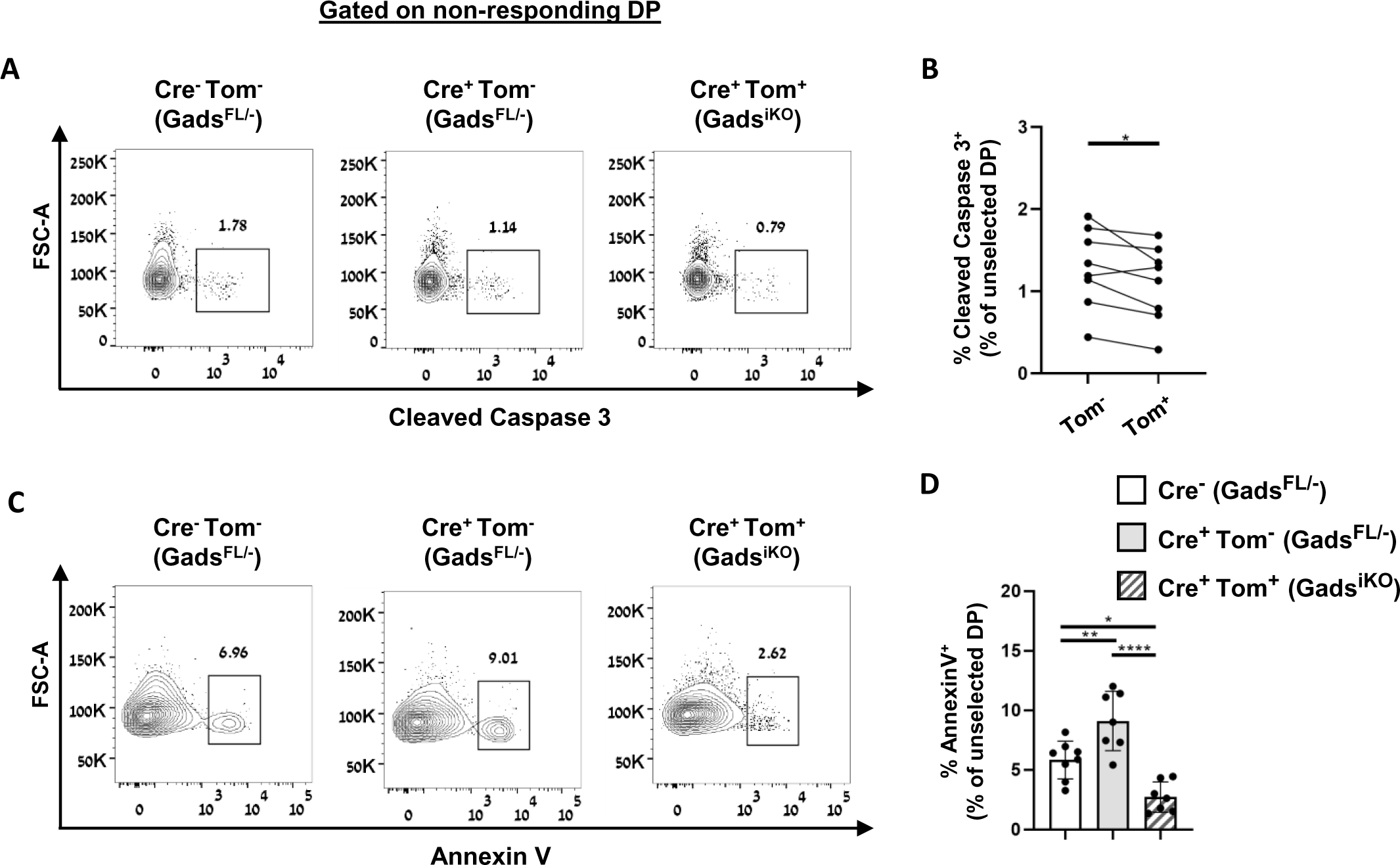
Gads facilitates death by neglect within the unselected DP population. **(A-D)** Detection of physiologic death by neglect. 2-2.5 weeks after the initiation of tamoxifen treatment, thymocytes were analyzed by FACS while gating on the live, non-responding DP population, as defined in Fig 4C. (**A**) Cleaved caspase 3 was detected by intracellular staining. (**B**) Paired comparison of the frequency of cleaved caspase 3^+^ cells, observed within Tom^+^ and Tom^−^ non-responding DP populations from 8 individual Cre^+^ mice. Statistical significance was determined by paired T-test. (**C**) Annexin 5 staining. (**D**) The frequency of annexin 5^+^ cells observed within the non-responding DP populations. n=8 Cre^−^ and 7 Cre^+^ mice. Statistical significance was determined by a one-way ANOVA Tukey’s multiple comparison test.

### Gads^iKO^ DP thymocytes exhibit reduced responsiveness in a model of experimentally-induced death by neglect

To explore whether the accumulation of non-responding Gads^iKO^ DP thymocytes can be attributed to a defect in death by neglect, we adapted a previously-published experimental approach, in which death of pre-selection CD5^lo^ DP thymocytes is induced upon CD8 cross-linking (44). This assay is based on the notion that binding of CD8 to class I MHC may serve to trigger death by neglect, if it occurs in the absence of TCR-mediated recognition of peptide-MHC.

For co-receptor cross linking, we incubated thymocytes at 37°C with avidin-beads that were either uncoated, or coated with different concentrations of CD8- or CD4-biotin antibodies. Annexin V and DAPI staining were then assessed, while gating on bead-bound, CD5^lo^ DP cells, as defined in Fig 7A.

**Fig 7.**
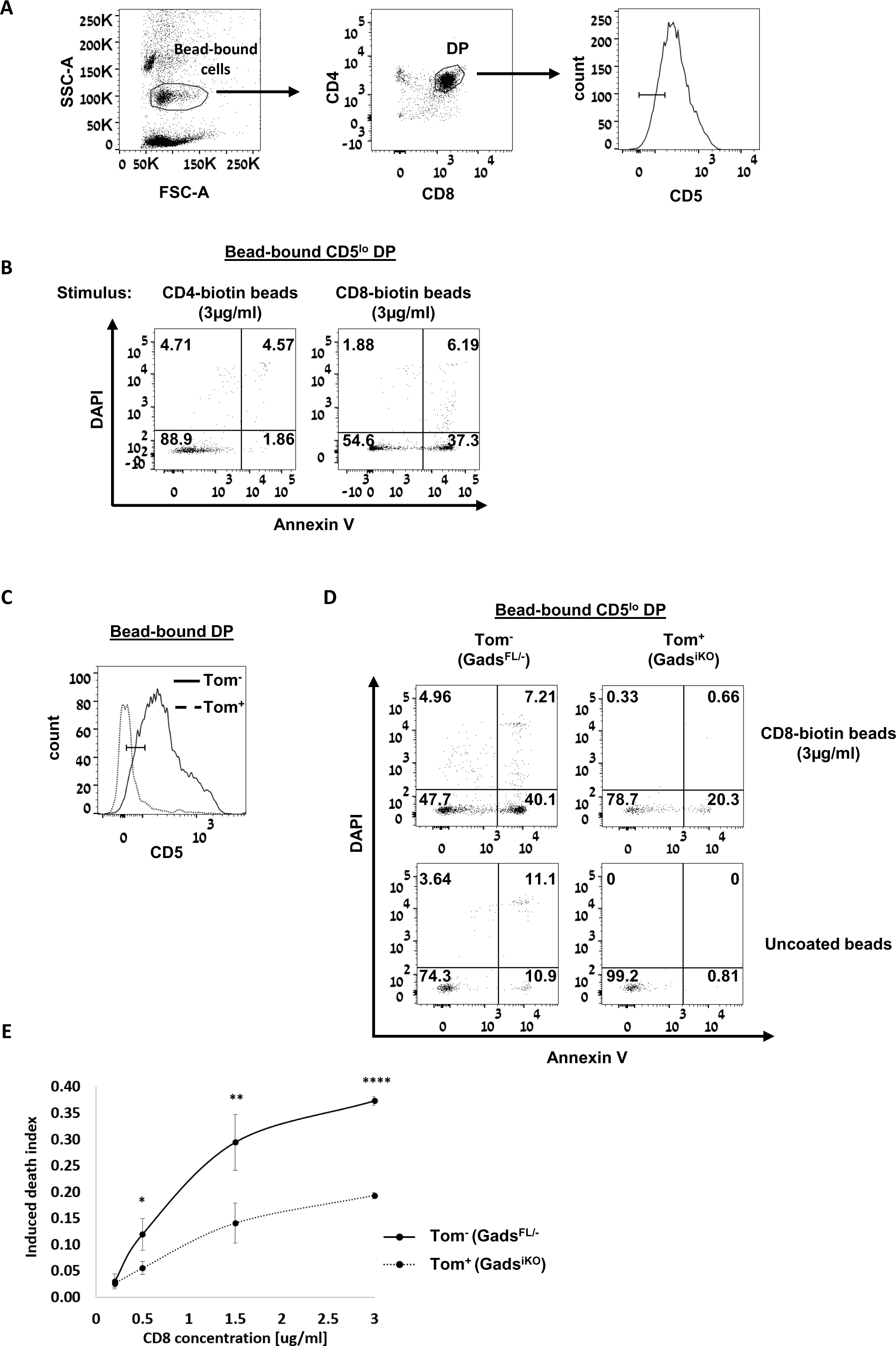
Gads^iKO^ DP thymocytes exhibit reduced responsiveness in a model of experimentally-induced death by neglect. Avidin-beads were pre-coated with CD4- or CD8-biotin, and then incubated for one hour with thymocytes, followed by staining with anti-CD4, -CD8, -CD5, Annexin V and DAPI. (**A-B**) Validation of experimental strategy, using wild-type (Cre^−^Tom^−^) thymocytes. (**A**) All results were assessed while gating on bead-bound CD5^lo^ DP thymocytes, using the gating strategy shown. (**B**) Representative DAPI and Annexin V staining within the bead-bound CD5^lo^ DP gate. (**C-E**) Experimentally induced death by neglect, in the presence and absence of Gads. Four weeks after the initiation of tamoxifen treatment, Tom^+^ and Tom^−^ thymocytes were mixed together and incubated with uncoated or CD8-biotin-coated beads. (**C**) The CD5^lo^ gate used in this experiment. (**D**) Representative DAPI and Annexin V staining within the Tom^−^ and Tom^+^ bead-bound CD5^lo^ gate. (**E**) Dose response of CD8-induced death by neglect, in the presence and absence of Gads. Beads were coated with 0, 0.2, 0.5, 1.5 or 3µg/ml CD8-biotin. The induced death index was calculated as described in Methods. n=4 technical repeats, error bars indicate standard deviation, similar results were obtained in at least two independent experiments. Statistically significant differences between Tom^+^ and Tom^−^ samples were determined using an unpaired T-test.

We chose a one hour time point to maximize the detection of early apoptotic cells (DAPI^−^ Annexin V^+^), while minimizing dead cells (DAPI^+^ Annexin V^+^). Under these conditions, CD8-biotin beads induced a marked increase in the frequency of early apoptotic cells, but no such increase was observed when thymocytes were incubated with CD4-biotin beads (Fig 7B).

To explore the role of Gads in this model of CD8-induced death, we incubated a mixture of Tom^+^ and Tom^−^ thymocytes with uncoated or CD8-biotin-coated beads, and analyzed the results while gating on a CD5^lo^ region within the bead-bound DP population (Fig 7C). Consistent with their lower rate of death by neglect *in vivo* (Fig 6C-D), Gads^iKO^ (Tom^+^) CD5^lo^DP thymocytes exhibited a lower frequency of early apoptotic cells following their incubation with control, uncoated beads (Fig 7D, lower panels). CD8 crosslinking induced a marked increase in the frequency of early apoptotic cells; however, the magnitude of this increase was reduced in the absence of Gads (Fig 7D). To determine the dose response of experimentally-induced early apoptosis, we performed the experiment using beads coated with increasing amounts of anti-CD8. (Fig 7E). The rate of induced death was significantly decreased in Gads^iKO^ thymocytes, suggesting that Gads plays an important role in this model of CD8-induced death by neglect.

Taken together, our data suggest two ways in which Gads regulates progression through the DP compartment. First, Gads is required for optimal responsiveness of DP thymocytes to self-peptide-MHC, as evidenced by the failure of Gads^iKO^ DP thymocytes to upregulate CD5 *in vivo*, and by their markedly reduced TCR responsiveness *ex vivo*. Second, Gads is required to promote death by neglect in CD5^lo^ DP thymocytes, both *in vivo* and in an *ex vivo* model of induced death. These twin defects likely explain the accumulation of CD5^−^ Gads^iKO^ DP thymocytes.

### Parallel progression of selected and unselected Gads^iKO^ thymocytes into the CD4 SP compartment

Despite the poor TCR-responsiveness of Gads^iKO^ DP thymocytes (see Fig 5C-D), Gads-deficient thymocytes progressed to the SP populations (see Fig 3A), raising the question of whether passage of Gads-deficient thymocytes into the SP compartments may be driven by positive selection. To address this question, we examined well-known surface markers of positive selection among SP thymocytes: increased expression of CD5, TCRβ, and CCR7, along with decreased expression of CD24 (45, 46). To exclude any potential non-SP populations from this analysis, we used “tight” SP gates, as illustrated in Fig 3A.

Consistent with positive selection, the vast majority of Gads-expressing CD4 SP thymocytes were found within a TCRβ^hi^CD5^hi^ gate. (Fig 8A, left panels). Moreover, most Gads^iKO^ (Tom^+^) CD4 SP thymocytes were found within the TCRβ^hi^CD5^hi^ gate (Fig 8A, right panel). Finally, nearly all CD8 SP thymocytes, whether Gads-expressing or Gads^iKO^, were found within the TCRβ^hi^CD5^hi^ gate (Fig 8B).

**Fig 8.**
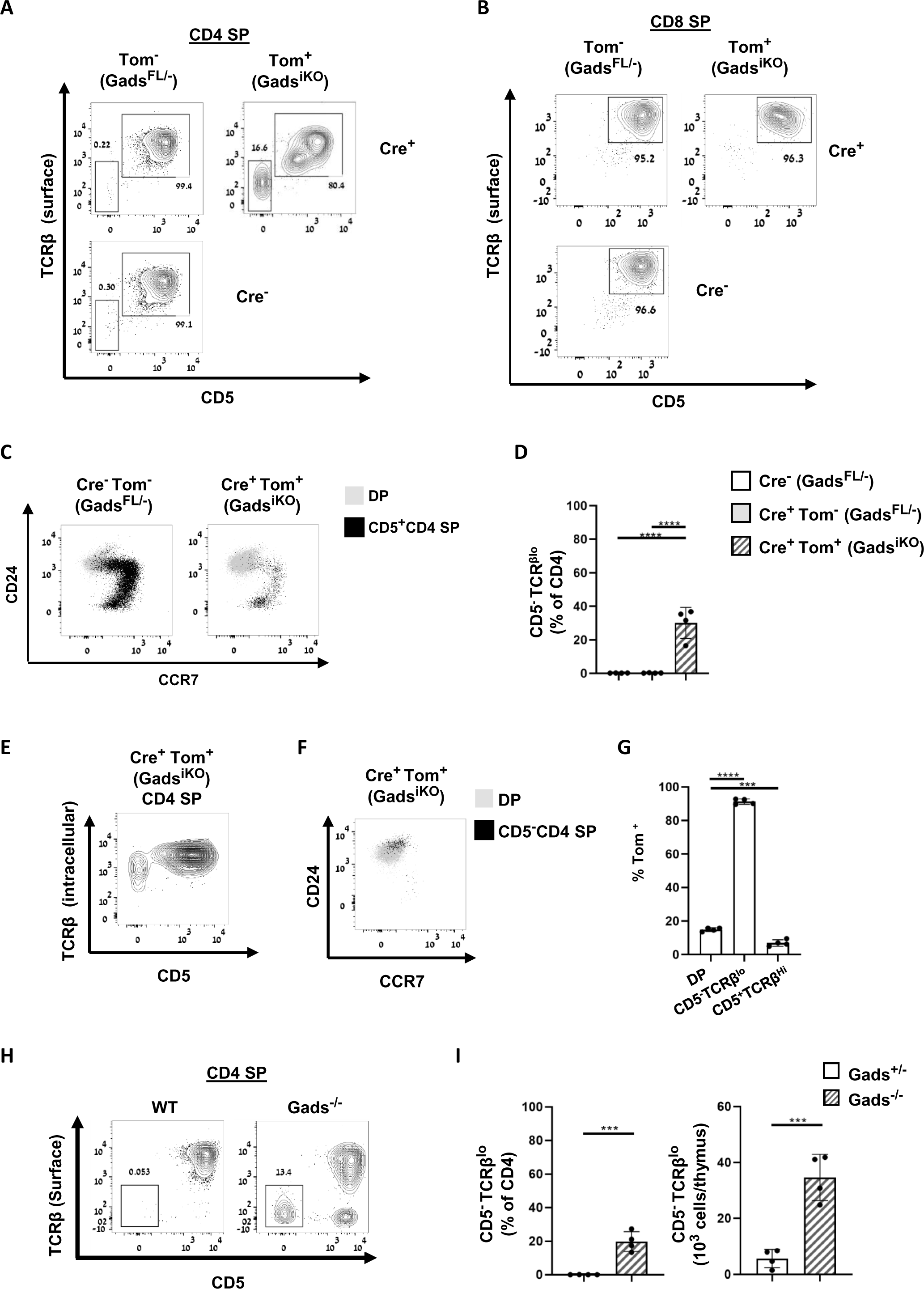
Parallel progression of selected and unselected Gads^iKO^ thymocytes into the CD4 SP compartment. (**A-G**) 2.5 weeks after the initiation of tamoxifen treatment thymocytes were analyzed by FACS while gating on the live CD4^+^ and CD8^+^SP population, defined as in Fig 3A. (**A and B**) Expression of CD5 and TCRβ within CD4 and CD8 SP populations was detected using surface staining. Representative result. (**C**) Expression of CCR7 and CD24 was detected using surface staining while gating on the indicated DP and CD5^+^CD4 SP populations. Representative result. (**D**) Quantitative analysis of the experiment shown in A. The frequency of CD5^−^ TCRβ^lo^ thymocytes within the CD4 SP population. Statistical significance was determined by one-way ANOVA Tukey’s multiple comparisons test. n= 4 mice of each genotype. (**E**) Intracellular TCRβ and surface CD5 expression in Gads^iKO^ CD4 SP thymocytes. (**F**) Expression of CD24 and CCR7 was detected as in C, but while gating on the Gads^iKO^ DP and CD5^−^ CD4 SP populations. (**G**) The percent of Tom^+^ thymocytes within each of the indicated gates. Statistical significance was determined by one-way ANOVA Dunnett’s multiple comparisons test. n= 4 Cre^+^ mice. (**H-I**) Thymocytes from 8.5-9 week old mice of WT and conventional KO mice were stained and analyzed by FACS. **(H)** Expression of TCRβ and CD5 was detected while gating on CD4 SP population using surface staining. Representative result. (**I**) The percent and the number of CD5^−^ TCRβ^lo^ defined in H was assessed. Statistical significance was determined by Unpaired T-test (n=4 WT, 4 KO mice).

Additional markers provided further support for the existence of positively-selected Gads^iKO^ SP populations. Whether Gads-expressing or Gads^iKO^, CD5^+^ CD4 SP thymocytes were found on a trajectory of increasing CCR7 and decreasing CD24, as compared to the DP population from which they derived (Fig 8C). These markers suggest that, despite low TCR-responsiveness in the DP compartment, a subset of Gads^iKO^ thymocytes may undergo TCR-driven positive selection.

A distinct population of Gads^iKO^ CD4 SP thymocytes was found in an unusual TCRβ^lo^CD5^−^ population, which was nearly absent in Gads-expressing cells (Fig 8A and D). This TCRβ^lo^CD5^−^ Gads^iKO^ CD4 SP population was observed at all time points that we examined from 2-6.5 weeks after the initiation of tamoxifen treatment (data not shown).

The TCRβ^lo^CD5^−^ CD4 SP population was clearly part of the T cell lineage, as demonstrated by its strong intracellular staining with TCRβ (Fig 8E). Nevertheless, cells in this population did not exhibit signs of positive selection. In addition to low expression of CD5, CD24 expression remained high and CCR7 remained low (Fig 8F). Together, these data suggest that thymocytes within the TCRβ^lo^CD5^−^ CD4 SP population passed into the CD4 SP compartment without undergoing positive selection.

To better understand the degree to which Gads affects positive selection, we examined the frequency of Tom^+^ cells in the selected (CD5^+^) and unselected (CD5^−^) CD4 SP populations, relative to their frequency in the DP population (Fig 8G). Consistent with an impairment of positive selection, the frequency of Tom^+^ cells was significantly reduced in the CD5^+^CD4 SP population. In contrast, the TCRβ^lo^CD5^−^ population was composed almost entirely of Tom^+^ cells, suggesting that the ability of unselected DP cells to progress to the SP compartment is selectively found in Gads-deficient cells.

A similar TCRβ^lo^CD5^−^ CD4 SP population was found in germline Gads-deficient mice (Fig 8H). Both the prevalence and the absolute number of TCRβ^lo^CD5^−^ CD4 SP thymocytes were markedly increased in germline Gads-deficient mice as compared to age-matched wild type mice (Fig 8I), Together, these results suggest that the TCRβ^lo^CD5^−^ CD4 SP population is an atypical population that is a conserved feature of Gads-deficient thymocyte development.

Taken together, our data suggest two potential pathways of CD4 SP development in the absence of Gads. On the one hand CD5^+^ Gads^iKO^ SP thymocytes appear to result from positive selection. On the other hand, CD5^−^ Gads^iKO^ appear to develop in the absence of positive selection; these thymocytes may derive from unselected DP thymocytes that did not undergo death by neglect; rather progressing as far as the CD4 SP compartment. We do not have any evidence that this population exits the thymus, and believe it may be a “dead end” thymic population that accumulates in the absence of Gads.

## Discussion

Despite years of research, the regulatory roles of Gads in thymic development remain unclear. To illuminate this issue, we developed a mouse model in which Gads-expressing and Gads-ablated thymocytes develop side by side in the same animal. Ablation was driven by a ubiquitously-expressed, tamoxifen inducible Cre recombinase, with tamoxifen administered 2-4 weeks prior to harvesting thymi. Thus, the Gads^iKO^ thymic populations that we analyzed were largely derived from progenitor cells that underwent ablation of Gads prior to entering the thymus. In this model, Gads-ablated thymocytes exhibited largely cell autonomous phenotypes that resembled germline Gads-deficient thymocytes, whereas Gads-expressing thymocytes in the same mouse phenocopied wild type thymocytes.

The co-development of Gads-expressing (Tom^−^) and Gads-ablated (Tom^+^) thymocytes allowed us to more rigorously identify developmental transitions at which Gads plays a role. Within the DN compartment, Gads was required for efficient β-selection, and for the consequent pre-TCR-induced increase in the expression of CD5.

Our results are consistent with the β-selection defect previously observed in germline Gads-deficient mice (38, 39), and with the failure of Gads-deficient DN thymocytes to respond to anti-CD3 stimulation (38). To gain further insight into this phenomenon, we developed an approach to assess physiologic pre-TCR signaling *in vivo*, based on the established relationship between pre-TCR signaling and CD5 expression (8). Our approach revealed that substantial populations of Gads^iKO^ DN thymocytes fail to upregulate CD5 or progress to DN4 despite expressing surface TCRβ. Among Gads^iKO^ cells that transited to DN4, cells with low to medium TCRβ expression failed to upregulate CD5, which was expressed only on DN4 thymocytes with the highest expression of TCRβ. These findings provide strong confirmation that the developmental defects of Gads-deficient DN thymocytes can be connected to a pre-TCR signaling defect.

Likewise, substantial evidence suggests that Gads is required in the DP compartment to support signaling through the mature αβ TCR. Previous studies demonstrated that Gads-deficient, TCR-transgenic mice have prominent defects in positive and negative selection (38, 47). We extended these studies to a polyclonal TCR repertoire by using CD5 as a physiologic marker that correlates with TCR signaling intensity *in vivo* (8). In both the inducible and the germline-deleted mouse model, most Gads-deficient DP thymocytes failed to upregulate CD5, indicating impaired responsiveness to self-MHC. Moreover, even CD5^+^ Gads^iKO^ DP thymocytes exhibited profoundly reduced TCR-induced calcium influx. Together, these results suggest a profound TCR-signaling defect in Gads-deficient DP.

The observed pre-TCR and TCR signaling defects likely reflect the role of Gads in mediating the TCR-induced recruitment of SLP-76 to LAT. This task is facilitated by the high affinity (low nanomolar) constitutive binding of Gads to SLP-76 (25). Grb2 most likely cannot compensate for Gads, since its binding to SLP-76 occurs at 1000-fold lower affinity (25). Thus, TCR-induced recruitment of SLP-76 to LAT is profoundly reduced in the absence of Gads, but is not affected by deletion of other Grb2-family members (32, 48).

Given the reduced TCR signaling competence of Gads-deficient DP thymocytes, one might expect to see increased death by neglect, which occurs in MHC-non-responsive wild-type thymocytes (10, 11). Contrary to this expectation, we report a striking accumulation of CD5^−^ DP thymocytes, which we observed both in the inducible and in the germline Gads-deficient models. This previously unreported phenotype first suggested to us that Gads may be required to induce the death of MHC-non-responding DP thymocytes. This notion is further supported by the reduced frequency of an annexin V^+^ cells among CD5^−^ Gads^iKO^ DP thymocytes.

Taken alone, the reduced frequency of death by neglect in situ is difficult to interpret. One possibility is that, in the absence of Gads, TCR signaling may be reduced to a level that is sufficient for survival but insufficient for efficient positive selection. Alternatively, our results may suggest a novel reinterpretation of the process of death by neglect. Rather that constituting a passive response to the absence of signaling events, the death of MHC-non-responsive DP thymocytes may be actively induced by particular signaling pathways that depend on Gads.

The notion that T cell-specific signaling pathways may be involved in triggering death by neglect was first suggested by Grebe *et al.*, who demonstrated that cross linking of CD8 induced apoptotic death, specifically in CD5^lo^, but not in CD5^hi^ DP thymocytes (44). This death could be prevented by co-crosslinking CD8 and CD3 (44). Based on these results Grebe *et al.* suggested that CD5^lo^ DP thymocytes are able to detecting signaling through CD8, and moreover, that in the absence of concomitant signaling through the TCR, the CD8-induced signaling pathway can trigger a form of death that may be related to the phenomenon of death by neglect. In support for their hypothesis, they note that early DP thymocytes express a nonsialylated form of CD8, that binds with high affinity to class I MHC (49, 50), and therefore may be able to induce signaling events in the absence of TCR ligation.

We recapitulated the observation of CD8-induced death of CD5^lo^ DP thymocytes, moreover, we extended this system to show that death is specifically induced by ligation CD8, but not CD4. Most importantly, we showed that Gads is required for optimal CD8-induced death in this experimental model.

To the best of our knowledge, our observation provides the first evidence that signaling elements downstream of the TCR may be involved in triggering the death of MHC-non-responsive DP thymocytes. Perhaps consistent with an involvement of Gads in cell death, Gads contains a caspase 3 cleavage site (51, 52); moreover, Gads-deficient T1 B cells were shown to be resistant to BCR-induced apoptosis (53).

Surprisingly, unselected, CD5^−^ thymocytes progressed as far as the CD4 compartment, where they remained in an immature, CD24^hi^TCRβ^lo^CCR7^lo^ state. This population likely corresponds to the CD24^hi^TCRβ^lo^ transitional CD4 SP population that accumulates in TCR-transgenic Gads-deficient mice (47). A population of CD24^hi^ CD8 SP thymocytes was likewise observed in cyclososporin A-treated HY-transgenic female mice (54). Together, these results suggest that upon partial disruption of TCR signaling, unselected thymocytes may advance to the SP compartments, but fail to fully mature. Whereas Dalheimer *et al.* interpreted their results as an indication of uncoupling of positive selection from developmental progression (47), we suggest that the developmental progression of unselected Gads-deficient thymocytes may reflect a failure of death by neglect.

In parallel, we observed distinct populations of positively-selected CD5^+^ SP thymocytes. The Gads-independent developmental progression of CD5^+^ populations suggests the possibility of Gads-independent TCR responsiveness. Consistent with this notion, several lines of evidence suggest that Gads is not essential for TCR responsiveness, but rather appears to regulate the probability of response to low intensity TCR signals (32, 39, 55). Residual Gads-independent TCR responsiveness may be sufficient to support the development of particular SP populations that require only low-level TCR signaling for their development. Alternatively, the TCR repertoire of positively-selected thymocytes may be skewed towards higher affinity receptors in the absence of Gads. We shall address the nature of the Gads-independent positively-selected populations in another manuscript.

Taken together, our results suggest a new paradigm in which Gads regulates the balance between positive selection and death by neglect. Both processes are reduced in the absence of Gads, resulting in the survival and developmental progression of unselected thymocytes.

## Acknowledgements

This research was supported by grants to DY from the Israel Science Foundation (283/22), the Rappaport Family Institute for Research in the Medical Sciences, the Minerva Center on Cell Intelligence and the Colleck Research Fund. The Biomedical Core Facility (BCF) of the Rappaport Faculty of Medicine provided access to FACS equipment, and BCF staff members Ofer Shenker, Amir Grau and Rotem Honen Kadosh provided excellent technical support. Gads-deficient mice were generously provided by C. Jane McGlade (University of Toronto). We thank our chief veterinarians Dr. Rona Shofti and Dr. Dvir Mintz, as well as Paul Zannou, and the rest of the excellent staff from the Technion Preclinical Authority for professional assistance with the care, treatment and housing of the mice. Pamela Schwartzberg and Dominic Golec from the NIH/NIAID and Leslie Berg from the University of Colorado provided much appreciated input into the interpretation of thymic phenotypes.

## Author contributions

RS and DY developed the study objectives and experimental strategy with key inputs from EH and MM. RS, EH and MM performed experiments, analyzed data and prepared the figures. NK designed and carried out all breeding and genotyping strategies for these experiments. This manuscript was written by DY and RS. All authors have read and agreed to the published version of the manuscript.

